# *doublesex* regulates sexually dimorphic beetle horn formation by integrating spatial and temporal developmental contexts in the Japanese rhinoceros beetle *Trypoxylus dichotomus*

**DOI:** 10.1101/328120

**Authors:** Shinichi Morita, Toshiya Ando, Akiteru Maeno, Takeshi Mizutani, Mutsuki Mase, Shuji Shigenobu, Teruyuki Niimi

**Affiliations:** Division of Evolutionary Developmental Biology, National Institute for Basic Biology, 38 Nishigonaka, Myodaiji, Okazaki 444-8585, Japan; Department of Basic Biology, School of Life Science, SOKENDAI (The Graduate University for Advanced Studies), 38 Nishigonaka, Myodaiji, Okazaki 444-8585, Japan; Mammalian Genetics Laboratory, Genetic Strains Research Center, National Institute of Genetics, Mishima, Shizuoka 411-8540, Japan.; Graduate School of Bioagricultural Sciences, Nagoya University, Chikusa, Nagoya 464-8601, Japan.; NIBB Core Research Facilities, National Institute for Basic Biology, 38 Nishigonaka Myodaiji, Okazaki 444-8585, Japan

## Abstract

Many scarab beetles have sexually dimorphic exaggerated horns that are an evolutionary novelty. Since the shape, number, size, and location of horns are highly diverged within Scarabaeidae, beetle horns are an attractive model for studying the evolution of sexually dimorphic and novel traits. In beetles including the Japanese rhinoceros beetle *Trypoxylus dichotomus*, the sex determination gene *doublesex* (*dsx*) plays a crucial role in sexually dimorphic horn formation during larval-pupal development. However, knowledge of when and how *dsx* drives the gene regulatory network (GRN) for horn formation to form sexually dimorphic horns during development remains elusive. To address this issue, we identified a *Trypoxylus*-ortholog of the sex determination gene, *transformer* (*tra*), that regulates sex-specific splicing of the *dsx* pre-mRNA, and whose loss of function results in sex transformation. By knocking down *tra* function at multiple developmental timepoints during larval-pupal development, we estimated the onset when the sex-specific GRN for horn formation is driven. In addition, we also revealed that *dsx* regulates different aspects of morphogenetic activities during the prepupal and pupal developmental stages to form appropriate morphologies of pupal head and thoracic horn primordia as well as those of adult horns. Based on these findings, we discuss the evolutionary developmental background of sexually dimorphic trait growth in horned beetles.

**Author Summary:**

Beetle horns are highly enriched in a particular family Scarabaeidae, although the shape, size and number of horns are diversified within the group. In addition, many scarab beetle horns are sexually dimorphic. It has been questioned how a particular group of beetles has originated and diversified evolutionary novel horns. Here we found the exact time when morphological sexual dimorphism of horn primordia appeared, estimated the onset of the developmental program for sexually dimorphic horn formation driven by Doublesex, and revealed that Doublesex regulates different aspects of cell activities of horn primordia depending on the spatiotemporal contexts. Our study provides our understanding regarding regulatory shifts in these mechanisms during the evolution of sexually dimorphic traits in horned beetles.

## Introduction

Beetle horns are used as weapons for intraspecific combats between males. Beetle horns display sexual dimorphism in many Scarab species, and their shapes, numbers, sizes and forming regions are highly diverged even among closely related species [1-3]. Interestingly, the diversified horn forms are associated with their fighting styles, such as scooping up, piercing and throwing [4]. Furthermore, beetle horns are thought to be an evolutionary novelty. They are an outgrowth structure derived not from an appendage but from a dorsal epidermal sheet. Elucidating how these novel traits were acquired in Scarab species will lead to better understanding the mechanisms of morphological diversification during evolution. Therefore, beetle horns are an attractive model for studying not only the association of trait novelty with sexually dimorphic development but also the evolution of novel traits.

Beetle horn development has been investigated in several horned beetles including the Japanese rhinoceros beetle, *Trypoxylus dichotomus* (Coleoptera, Scarabaeidae, Dynastinae). In *T. dichotomus*, male adults have sexually dimorphic exaggerated horns on the head and prothorax, which are used in combats among conspecific males as weapons [5,6]. The head horn is shaped like a plow with a long stalk, and bifurcated twice at the distal tip, while the prothoracic horn is shorter than the head horn, and bifurcated once at the distal tip. During development, the horns are first formed as thickened epidermal primordia at the prepupal stage. Male adults have exaggerated horns at the head and prothoracic regions, whereas females do not have these structures. However, females have a small head horn at the pupal stage (Fig. 1B, Fig. S1). Sexual dimorphism of horns can be first observed at the prepupal stage [7,8]. In males, the prepupal horn primordia are elongated immediately after pupation, to the same length as an adult horn [9]. However, the shape of a pupal horn is rounded, and slightly larger than an adult horn (Fig. 1B, Fig. S1). In horned beetles, the pupal horn is reported to be remodeled during the pupal period to form the final adult horn [7,10,11].

**Fig. 1.**
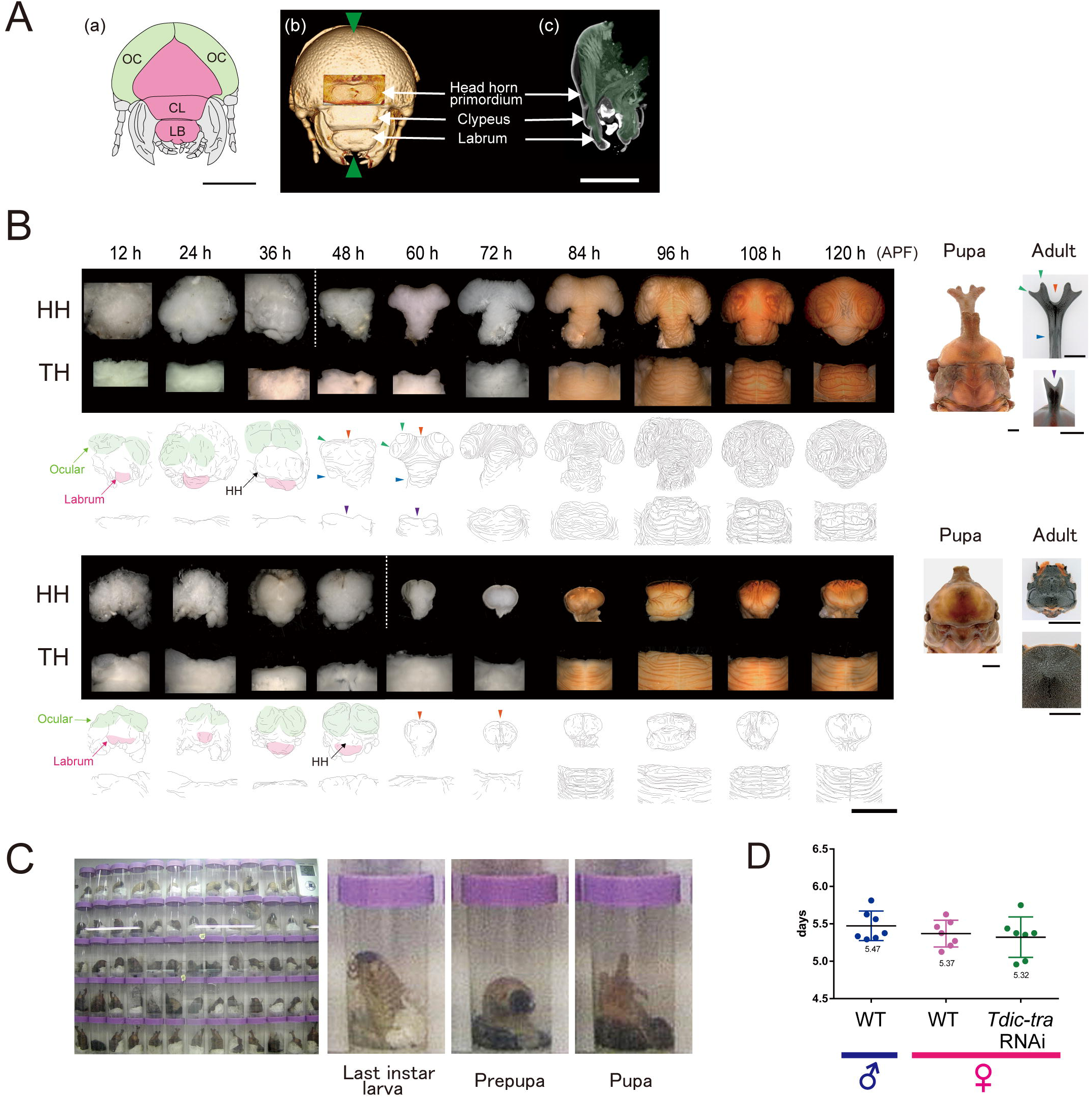
Morphological change of horn primordia and the prepupal period in *Trypoxylus dichotomus.* (A) (a) Schematic of the frontal view of larval head from *T. dichotomus*. Green, ocular (OC); Magenta, clypeolabrum; CL, clypeus; LB, labrum. (b), (c) Micro-CT images of prepupal head at 24 h APF from *Trypoxylus*. (b) The frontal view. Green arrowheads indicate the position of a sagittal section in (c). (c) The sagittal section of head. Head horn primordia were formed above the clypeus in the clypeolabrum (Movie 1a). (B) Morphological changes of head and thoracic horn primordia during larval-pupal development revealed by the time-lapse photography system. Both in males (the upper panels in the upper half) and in females (the upper panels in the lower half), horn primordia were formed in the clypeolabrum. In males, the frontal views are presented concerning the head horn primordia from 12 h APF to 36 h APF, and the anterodorsal views are presented concerning the head horn primordia from 48 h APF to 120 h APF. In females, the frontal views are presented demonstrated the head horn primordia from 12 h APF to 48 h APF, and the anterodorsal views are presented demonstrating the head horn primordia from 60 h APF to 120 h APF. The dorsal views are presented demonstrating thoracic horn primordia. APF, after pupal-chamber formation; HH, head horn primordia; TH, thoracic horn primordia; Orange arrowheads, the firstly bifurcated distal tips of a head horn; Green arrowheads, the secondly bifurcated distal tips of a head horn; Blue arrowheads, the long column shaped stalk of a head horn; Purple arrowheads, the bifurcated distal tips of a thoracic horn. (C) Snapshots of a last-instar larva, prepupa and pupa under a time-lapse photography (Movie 2). (D) The durations of the prepupal periods in males and females estimated using the time-lapse photography system. Median, the upper/lower quantile, and minimum/maximum values were presented by a box‐ and whisker‐ plot. The median duration of prepupal periods in males and females were 131 hours and 129 hours, respectively. The mean duration of prepupal period in *Tdic-tra* RNAi females was 127 hours, which was slightly shorter (4 hours) than that of normally developed females.

In insects, orthologs of a transcription factor gene *doublesex* (*dsx*) play a pivotal role in sexual differentiation, and are involved in formation of sexually dimorphic traits by expressing sex-specific isoforms (*dsxM* and *dsxF*) [8, 12-17]. In addition, tissue-specific higher expression of *dsx* is detected where sexually dimorphic structures are formed to induce their development [17-21]. In horned beetles including *T. dichotomus*, the *dsx* orthologs also regulate sexually dimorphic horn formation during larval-pupal development. RNA interference (RNAi) targeting *dsx* results in intersexual phenotypes both in males and females [8,14].

Molecular mechanisms of sex-specific splicing of *dsx* has been intensely studied in *Drosophila melanogaster*, and the sex determination cascade plays a pivotal role [12, 22]. In *D. melanogaster*, *Sex-lethal* (*Sxl*), the master sex determination gene, initiates the sex determination cascade. *Sxl* encodes an RNA-binding protein that directly binds to target RNAs. Functional Sxl protein is translated only in females [23-25], and controls sex-specific splicing of *transformer* (*tra*) to produce functional Tra protein only in females [26]. Then, the Tra molecule specifically expressed in females forms a heterodimer with an ubiquitously expressed RNA-binding protein, Transformer2 (Tra2), to regulate the sex-specific alternative splicing of *dsx* by directly binding to *dsx* transcripts [27,28]. Resultant *dsxF* isoform in females and *dsxM* isoform in males regulate a battery of downstream genes to form sexually dimorphic traits by binding to the target DNA sequences with the Dsx binding motif [29,30], and by activating or repressing their transcription [31]. Intersex (Ix) functions as a female-specific co-activator by directly binding to female-specific isoforms of the DsxF protein, and regulates development of female-specific traits [32].

The above *Sxl*-[*tra/tra2*]-[*dsxF/ix*] pathway is turned on only in females. In males, default splicing results in expression of the male-specific splicing variant, *dsxM* [12,22]. Loss of function of the sex determination genes in females, and gain of function of those in males result in sex transformation [12,22].

Orthologous genes corresponding to the *D. melanogaster* sex determination genes described above are conserved among many holometabolous insects. However, whether these genes are involved in sex determination pathways to form sexually dimorphic horns remains elusive. In addition, when and how *dsx* interacts with the gene regulatory network (GRN) for horn formation to drive cellular activities such as cell growth, cell death and cell movement to form sexually dimorphic is also unknown.

To understand the developmental and genetic mechanisms underlying the sexually dimorphic horn formation in *T. dichotomus*, we first described a precise time course of the morphogenetic changes of male and female horn primordia during larval-pupal development using time-lapse photography. Next, we examined the function of putative sex determination genes in *T. dichotomus* using larval RNAi by focusing on orthologs of known *D. melanogaster* sex determination genes. Moreover, we investigated the initiation timing of GRN for horn formation by knocking down *Tdic-tra* at multiple developmental timepoints, and by evaluating the extent of sex transformation phenotypes. Based on these experiment, we concluded that the GRN for horn formation, which is supposed to be modified by *dsxM* and *dsxF* functions in males and females, is driven at a very early stage of larval-pupal development before clear morphological changes in horn primordia can be detected. Furthermore, we show that *dsxM* has different functions in both the prepupal and pupal stages during the formation of appropriate morphologies in pupal and adult horns in males. Based on these findings, we discuss the evolutionary developmental background of sexually dimorphic horn formation in horned beetles.

## Results

### Development of sexual dimorphism in *T. dichotomus* horn primordia

To identify the developmental timepoint when sexual dimorphism of horns first appears in *T. dichotomus*, we described morphological changes of head and thoracic horn primordia during the prepupal period. Micro-CT analysis of a head horn primordium at an early stage (24 h APF) in the prepupa revealed that a head horn primordium was formed in the clypeolabral region during the prepupal stage as investigated in *Onthophagus taurus*, *Onthophagus sagittarius* and *Onthophagus gazella* (Fig. 1A, B, Movie 1a, b) [3,33,34]. In addition, head horn primordia were formed above the clypeus in the clypeolabrum (Fig. 1A, Movie 1a, b). To determine the exact timepoint when protrusion of the primordium is initiated, we established a time-lapse photography system (Fig. 1C). Until recently, developmental staging of *T. dichotomus* prepupae had been difficult because they form pupation chambers underground and pupate in them. Using our time-lapse photography system, we found that the head-rocking behavior at the end of pupal-chamber formation can be an unambiguous marker for the initiation of the prepupal stage (Movie 2). We could minimize the developmental deviation between individuals within 12 hours using this precise marker (Fig 1C, Movie 2). The average prepupal period was 5.5 ± 0.19 days (131 ± 4.7 hours) in males and 5.4 ± 0.17 days (129 ± 4.3 hours) in females (Fig. 1D). Based on this staging paradigm, we manually dissected out horn primordia every 12 h after pupal-chamber formation (APF). We found that sexual dimorphism of horn primordia appeared at 36 h APF (Fig. 1B). Therefore, we concluded that the GRN driving horn sexual dimorphism formation would be activated before 36 h APF in *T. dichotomus*. In addition, we also found that apolysis occurring at 36 h APF can be another ununbiguous developmental marker. Larval mandibular tendons tightly connecting mandibular muscles and apodemes (Movie 1b) [35,36] were completely detached from apdemes at

36 h APF along with the apolysis occurring at every body part including the neighboring ocular region to know the timing of sexual dimorphism formation.

Identification of genes regulating the sex-specific splicing of *Tdic-dsx* during *T. dichotomus* horn formation In *T. dichotomus*, regulatory factors associated with sex-specific splicing of *dsx* had not been identified. We searched for such regulatory factors focusing on *T. dichotomus* orthologs of *D. melanogaster* sex determination genes (*Sxl*, *tra*, *tra2* and *ix*) [12,22].

First, we investigated whether these genes produce sex-specific splicing variants in *T. dichotomus* by RT-PCR. All of these genes were expressed in male and female prepupal head horns (Fig. 2A light green bars, Fig. 2B, Table S1). Among these genes, sex-specific splicing variants were detected only in *Tdic-tra* (Fig. 2B). Next, to test whether these genes function as sex determination genes, we performed larval RNAi experiments using dsRNA targeting the common regions between sexes (Fig. 2A, black bars, Table S1, Table S2) [8].

**Fig. 2.**
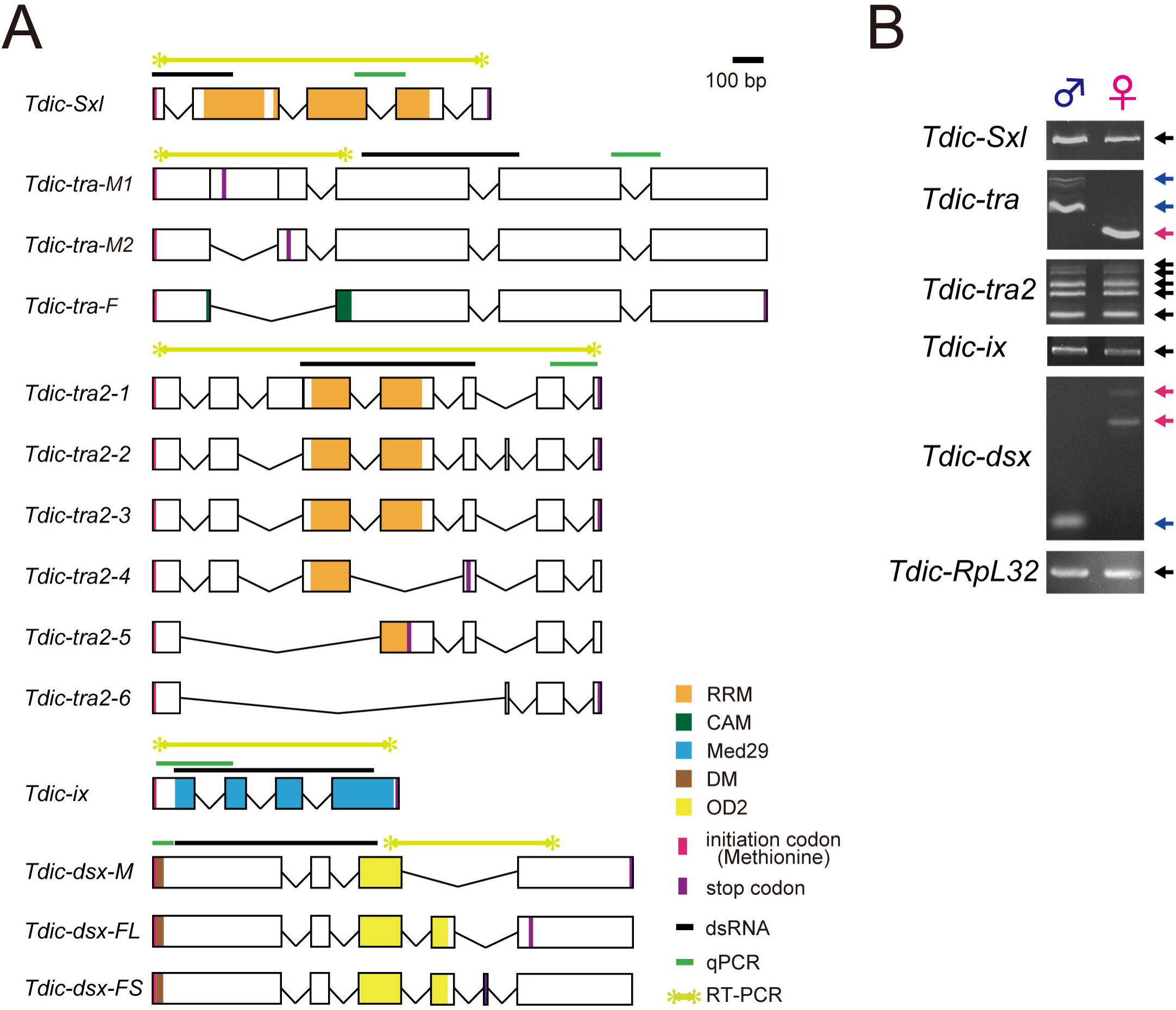
Schematic representation of putative exon–intron structures and sex-specific splicing patterns of the sex determination genes. (A) Schematics of putative exon-intron structures of the sex determination genes in *T. dichotomus*. Orange box, RRM (RNA binding motif); Green box, CAM (C, *Ceratitis*; A, *Apis*; M, *Musca*) domain [38,49], which is conserved in all *tra* orthologs identified in other insects except for those in *Drosophila* species and the sandfly (*Phlebotomus papatasi*) [48]; Light blue box, Med29 (mediator complex subunit 29) domain; Brown box, DM (Doublesex/Mab-3 DNA-binding) domain; Yellow box, OD2 (Oligomerization dmain 2); Black bars, template sequences for dsRNA synthesis; Green bars, the amplified regions in qRT-PCR analysis; Light green bars, the amplified region to test sex-specific splicing variants; Magenta lines, translation start sites; Purple lines, stop codons. (B) Evaluation of expression and sex-specific splicing patterns in the sex determination genes in male and female prepupal head horn primordia. Template cDNAs were derived from prepupal horn primordia at 72 h APF. *Tdic-RpL32* was used as an internal control for RT-PCR. Blue arrows, PCR products with male specific splicing variants; Magenta arrows, PCR products with female specific splicing variants; Black arrows, PCR products amplified in both sexes.

In females, the morphological sex transformation was observed in the RNAi treatments targeting *Tdic-tra* and *Tdic-ix*, whereas no morphological changes were observed in males (Fig. 3A). Such female-specific sex transformation phenotypes were comparable with the mutant phenotypes of *tra, tra2* and *ix* in *D. melanogaster* [12,22]. Concerning *Tdic-tra2,* we could not observe its adult phenotype because of the prepupal lethality in both sexes. The phenotypes in *Tdic-tra* RNAi females and *Tdic-ix* RNAi females were different and the *Tdic-ix* RNAi phenotype was similar to the effects of the *Tdic-dsx* RNAi phenotype (Fig. 3A). In the *Tdic-tra* RNAi females, ectopic horn formation was observed both the head and prothorax (Fig. 3A), whereas in *Tdic-ix* RNAi females, ectopic horns were formed only in the head, and these horns were significantly shorter than the ectopic head horns in *Tdic-tra* RNAi females (Fig. 3A, B). Such a difference in morphology was also observed in another sexually dimorphic structure, the intercoxal process of prosternum (IPP). Male IPPs are generally larger than female IPPs (Fig. 3A). Although both the *Tdic-tra* and *Tdic-ix* RNAi females have larger IPPs than female controls (*EGFP* RNAi), *Tdic-tra* RNAi females have much larger IPPs than *Tdic-ix* RNAi females (Fig. 3A). These results and the intermolecular interactions reported in *D. melanogaster* led us to predict that *Tdic-tra* may also regulate the sex-specific splicing of *Tdic-dsx* in *T. dichotomus*. To test this, we investigated the splicing patterns of *Tdic-dsx* in the above RNAi-treated males and females. We designed PCR primer sets to amplify the region including the whole female specific exon, which is spliced out in males (Fig. 2A, light green bars, Table S1). We found that the sex-specific splicing pattern observed in wild type (Fig. 2B) was switched in *Tdic-tra* and *Tdic-tra2* RNAi treatments, whereas splicing patterns were not changed by *Tdic-Sxl* and *Tdic-ix RNA*i treatments (Fig. 3C). These data indicate that *Tdic-tra* and *Tdic-tra2* regulate female-specific splicing of *Tdic-dsx* in *T. dichotomus*. The switching of sex-specific splicing of *Tdic-dsx* and the resultant morphological sex transformation in *Tdic-tra* RNAi females implies that the sex-specific splicing regulation of *Tdic-dsx* by *Tdic-tra* is also conserved in *T. dichotomus* as in other holometabolous insects investigated so far (*D. melanogaster*; [37], the housefly *Musca domestica*; [38], the Mediterranean fruit fly *Ceratitis capitata*; [39], the Australian sheep blowfly *Lucilia cuprina*; [40], the parasitic wasp *Nasonia vitripennis*; [41], the oriental fruit fly *Bactrocera dorsalis*; [42] and the red flour beetle*, Tribolium castaneum*; [43,44]). Taken together, we concluded that *Tdic-tra* functions as a sex determination gene during horn formation in *T. dichotomus*.

**Fig. 3.**
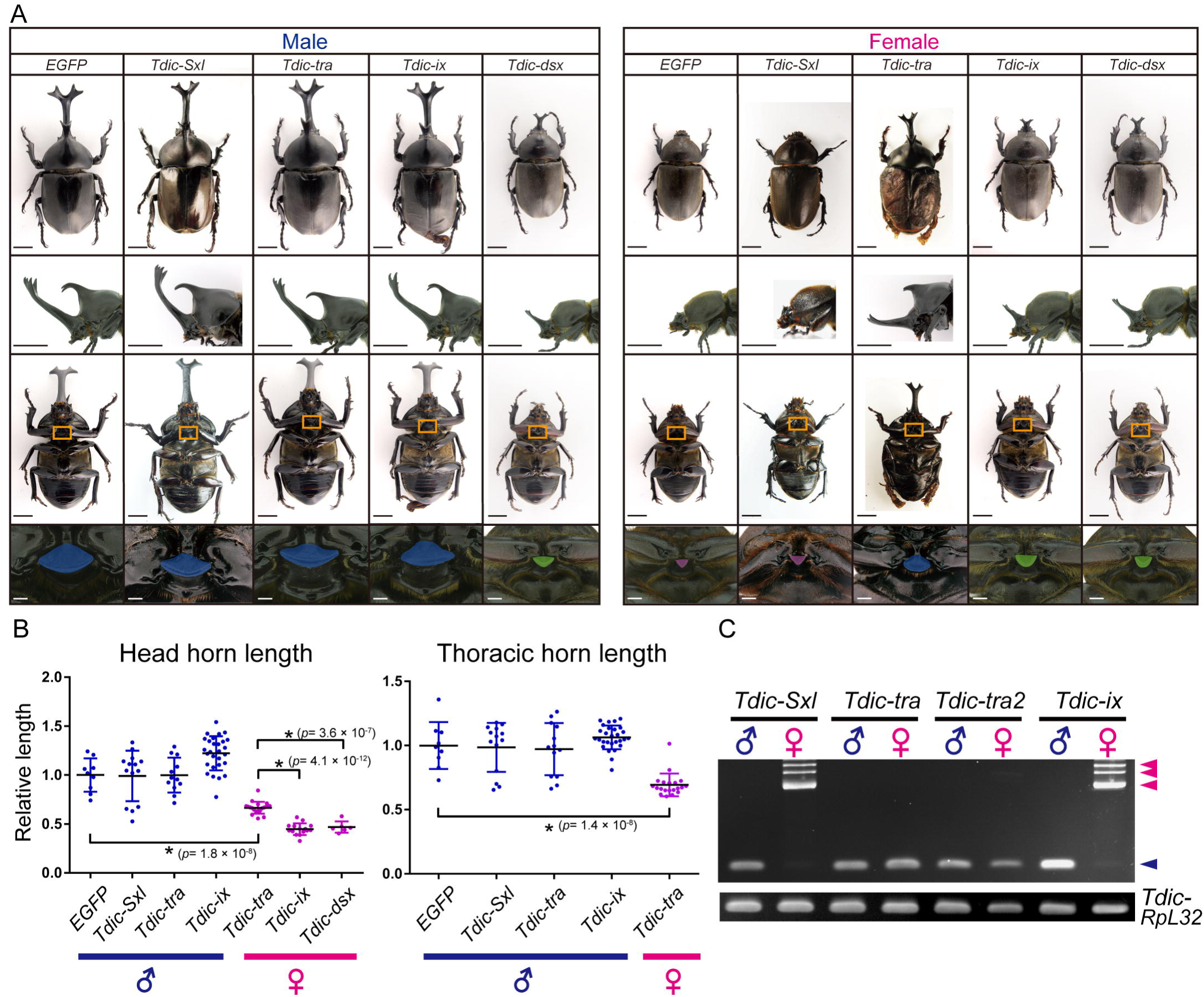
RNAi-mediated loss-of-function phenotypes of the sex determination genes. (A) Representative individuals in each RNAi treatment in males (the left half) and females (the right half). Each dsRNA was injected into last-instar larvae. The negative control RNAi treatment (*EGFP* dsRNA) showed no morphological defects. The upper row, the dorsal views of adults; the second row, the lateral views of a head and a prothorax in adults; the third row, the ventral views of adults; the forth row, magnified views of orange squares (IPP, intercoxal process of prosternum) in the third row. Scale bars are 1 cm in the upper three rows and 5 mm in the fourth row. (B) Quantification of the relative head and thoracic horn length in RNAi-treated individuals. Relative head and thoracic horn length in RNAi-treated males and females are plotted in the blue dots and the magenta dots, respectively. The relative horn lengths were standardized by dividing the horn length by the body size in each RNAi treated individual and by the mean horn length of the *EGFP* RNAi-treated males. Differences in horn length among individual RNAi treatments was compared using student’s t-test with Bonferroni correction. Asterisks denote significance: n.s., p > 0.05; *, p < 0.05. (C) Sex-specific splicing of *Tdic-dsx* in RNAi treatments targeting *Tdic-Sxl, Tdic-tra, Tdic-tra2* and *Tdic-ix*. Blue arrowheads, male specific splicing patterns. Magenta arrowheads, female specific splicing patterns. *Tdic-RpL32* was used as an internal control for RT-PCR

Estimation of the onset of the developmental program for sexually dimorphic horn formation using *Tdic-tra* RNAi

As sexual dimorphism of horn primordia appeared at 36 h APF (Fig. 1B), we speculated that the onset of developmental program for sexually dimorphic horn formation is driven before 36 h APF. To estimate this timepoint more accurately, we performed *Tdic-tra* RNAi in females at multiple developmental timepoints during pupal chamber formation periods and prepupal periods, and evaluated the extent of sexual transformation in horns.

If the timing of *Tdic-tra* RNAi in females is early enough, the ectopic *Tdic-dsxM* would be expressed in female horn primordium from the onset of developmental program for sexual dimorphism formation, and full sexual transformation can be achieved. On the other hand, the later the timing of *Tdic-tra* RNAi treatment becomes, the more the initial phases of the male-specific horn formation program driven by ectopically expressed *Tdic-dsxM* are trimmed, and at the same time repressed by normally expressed *dsxF*. Therefore, by determining the latest RNAi injection timing when a full sexual transformation phenotype is observed, we can estimate the onset of the sexually dimorphic horn formation program mediated by the ectopic *Tdic-dsxM* expression.

In this experiment, fully matured female last instar larvae and unstaged female prepupae were injected with *EGFP* or *Tdic-tra* dsRNA (Table S3). We performed time-lapse photography until pupation using these larvae, and retrospectively estimated the exact timing of injection before pupation. *Tdic-tra* RNAi females treated later than the timepoint at 130 hours before pupation showed either no morphological changes or incomplete sex transformation of head or horn, if any (Fig. 4A(ii) and (iii), B green dot). On the other hand, *Tdic-tra* RNAi females treated earlier than the timepoint at 134 hours before pupation formed fully developed male horns (Fig. 4A(i), B magenta dot). In this experiment, the prepupal period of wild type females and *Tdic-tra* RNAi females were comparable (Fig. 1D). Therefore, these RNAi experiments suggested that the estimated timepoint of the onset of the developmental program for sexually dimorphic horn formation is around 134 hours before pupation.

**Fig. 4.**
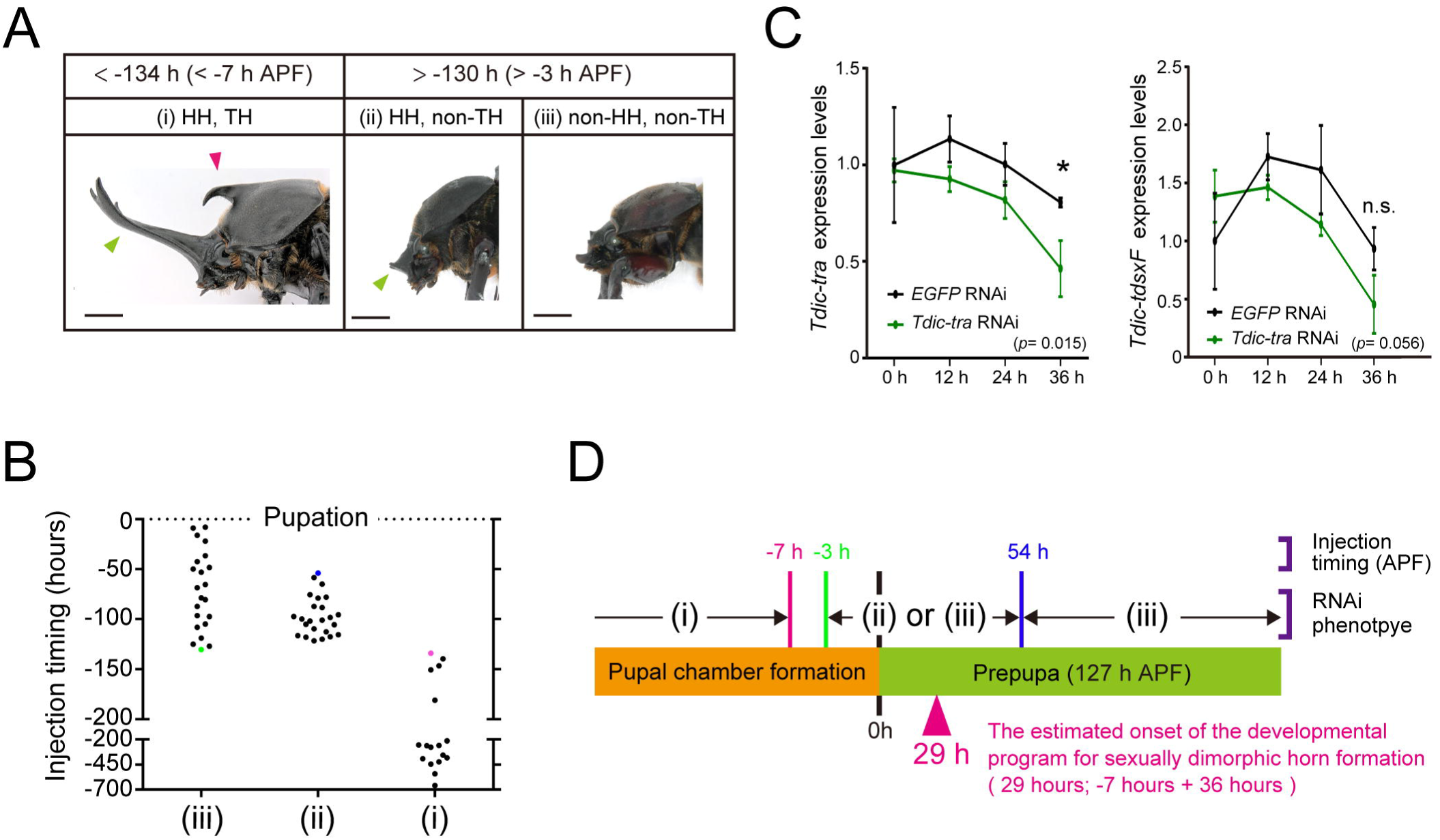
Estimation of the onset of the *Tdic* - *dsx* dependent horn developmental program by *Tdic-tra* RNAi. (A) Relationship between the timepoints of *Tdic-tra* dsRNA injection and the extent of sex transformation. The extent of the sex transformation (masculinization) phenotypes of *Tdic-tra* RNAi females were categorized into three classes: (i) prominent HH and TH were formed (< −134 h APF); (ii) only small HH was formed (−130 h APF ∼ 54 h APF); (iii) neither the HH nor TH was formed as in normal females (> −130 h APF). HH, head horn primordia; TH, thoracic horn primordia. (B) Timepoints of *Tdic-tra* dsRNA injection before pupation in each class were plotted. The green dot, the pink dot, and the blue dot indicate 130 h APF, 134 h APF, and 54 h APF, respectively. (C) Time course of relative *Tdic-tra* and *Tdic-dsxF* mRNA expression level after *Tdic-tra* dsRNA injection at 24 h APF. mRNA expression levels at each timepoint were quantified by qRT-PCR. The expression levels of *Tdic-tra* were significantly decreased at 36 hours after injection. Asterisks denote significance: n.s., p > 0.05; *, p < 0.05. (D) Summary of sex transformation phenotypes and the estimated onset of the *Tdic-dsx* dependent horn developmental program. The magenta, green and blue lines are the boundary timepoints, which divide developmental stages in each of which the same phenotypic category was observed. Magenta arrowheads indicate the timepoints corrected by the mRNA degradation delay (36 hours), which estimated by qRT-PCR. 0 h, onset of prepupa; APF, after puapal-chamber formation; Magenta arrowhead, the estimated onset of the developmental program for sexually dimorphic horn formation driven by *Tdic-dsx*.

Since this timepoint was estimated by means of RNAi, we speculated that there should be time lag between the timing of injection and the timing of the decrease in functional protein levels followed by the mRNA degradation. Thus, we also quantified expression dynamics of *Tdic-tra* mRNA after RNAi treatment by qRT-PCR. It was technically impossible to monitor the expression dynamics of *Tdic*-*dsx* at 134 hours before pupation, which is estimated to be 7 hours before pupal chamber formation (127 minus 134) (Fig. 1D, Fig. 4D). Then, we monitor the expression dynamics as early as possible (dsRNA injection at 24 h APF), instead. The expression levels of *Tdic-tra* and *Tdic-dsxF* were quantified every 12 hours up to 36 hours after injection (Fig. 2A, green bars, Fig. 4C). As a result, the expression levels of *Tdic-tra* and *Tdic-dsxF* were significantly decreased at 36 hours after injection (Fig. 4C). Since mRNA started to be degraded 36 hours after RNAi treatments (Fig. 4C), we estimated that this timepoint corresponded to 29 hours APF (−7 plus 36) (Fig. 4D). Taking morphological data into account, this timepoint corresponded to 7 hours before the initial sexual dimorphism appears in horn primordia (Fig.1B, Fig. 4D).

### Sexually dimorphic morphogenesis in male and female horn primordia

*T. dichotomus* male adults have exaggerated horns at the head and prothoracic regions whereas female adults only have three small protrusions at the clypeolabral region (Fig. 5A, magenta arrowhead). We wondered whether the male head horn and three female head protrusions were homologous or not. Since a head horn primordium is formed in both males and females, and occupies the almost entire clypeolabral region in head during prepupal stages, head horn primordiua formed in males and females seemed to be homologous structures (Fig. 1B). However, due to the lack of clear morphological landmarks indicating homology between male head horn and female head protrusion, their homology remained elusive. Unexpectedly, however, we obtained an intermediate sexual transformation phenotype of *Tdic-tra* RNAi to solve this problem. When we injected *Tdic-tra* dsRNA in small amounts (2.5 μg), we obtained several adults possessing both the three small protrusions similar to those in females, and a small ectopic anterior protrusion, seemingly analogous to a male head horn (Fig. 5B, C). This result suggests that at least the anterior region of a male head horn is not homologous to female head protrusions.

**Fig. 5.**
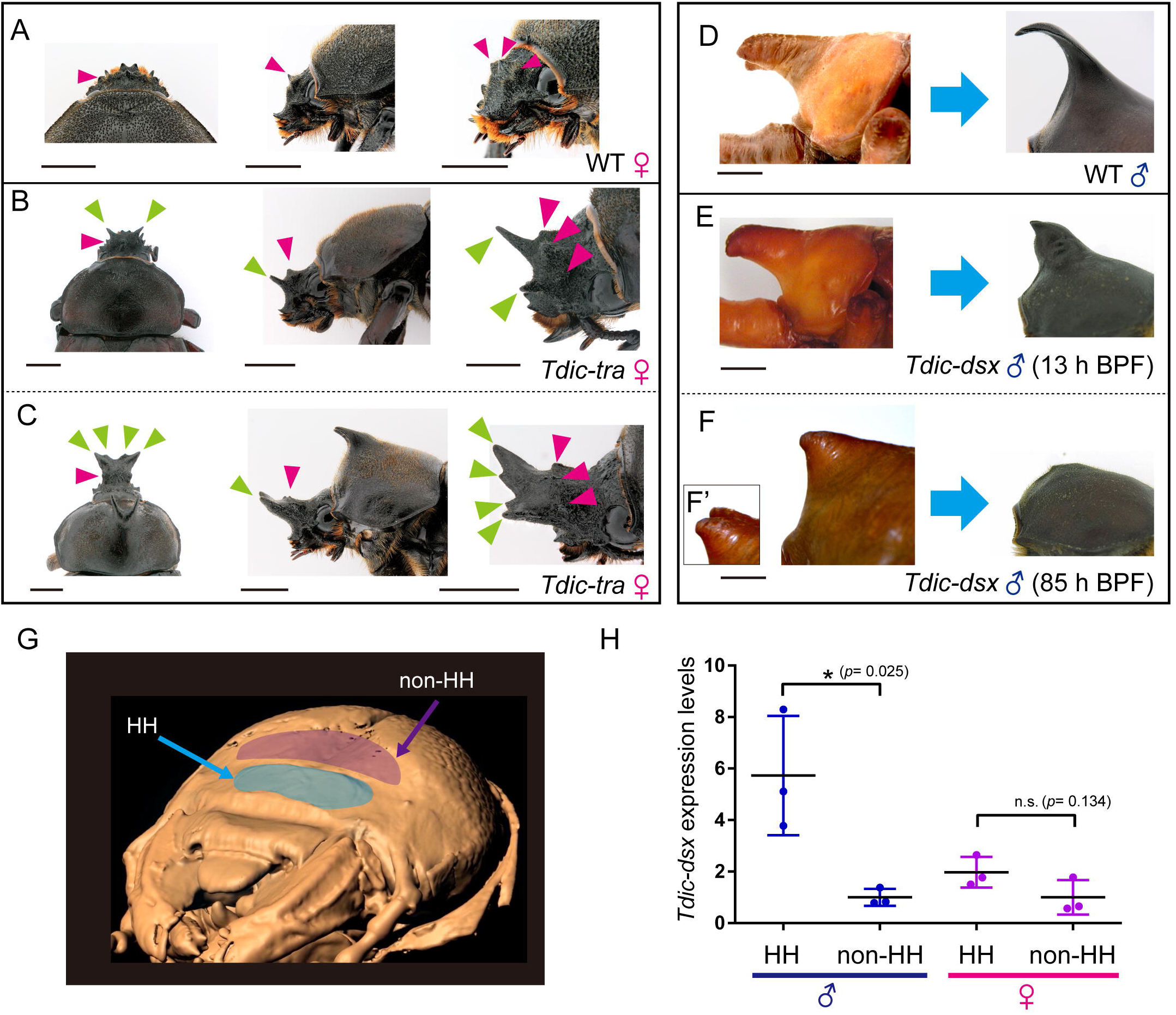
Horn formation phenotypes induced by *Tdic-tra* or *Tdic-dsx* RNAi at late prepupal stages. (A-C) Comparison of wild type female head and ectopic intermediate sexual transformation head horn in female heads induced by *Tdic-tra* RNAi treatments. (A) A wild type female head. (B) Small ectopic head horn formation in a female head. (C) Middle-sized ectopic head horn formation in a female head. Three small protrusions are formed in clypeolabrum on the wild type female head (A-C, magenta arrowheads), whereas ectopic small‐ or middle-sized ectopic head horns were formed in the region anterior to the three small protrusions (B, C, green arrowheads) in *Tdic-tra* RNAi treatments. (D-F) Thoracic horn remodeling during pupal-adult development in males. (D) A wild type male thoracic horn. (E) Unremodelled thoracic horn formed by a late *Tdic-dsx* RNAi treatment (13 BPF). (F) Almost complete reduction of a thoracic horn with an early *Tdic-dsx* RNAi treatment (85 BPF). (F’) dorsolateral view. (G) The epidermal regions used for qRT-PCR. The 3D volume image was reconstructed from sequential micro CT images. HH, head horn primordium; Blue, head horn primordium (HH); Purple, the head epidermis (non-HH). (H) The relative *Tdic-dsx* expression level in head horn primordia and head epidermis at 36 h APF in both sexes. Asterisks denote significance: n.s., p > 0.05; *, p < 0.05. BPF, before pupal-chamber formation; HH, head horn primordium epidermis; non-HH, the head epidermis.

The male and female horn primordia were formed above the clypeus in clypeolabrum during larval-pupal development (Fig. 1A, B). However, the adult male head horn and adult female protrusions are formed from different regions in a clypeolabrum (Fig. 5A-C). The shape of pupal horn is rounded, and slightly larger than that of male adult horns. The male and female pupal epithelia in the head horn primordia are remodeled during the pupal stage to form respective adult head structures (Fig. 5D, Fig. S1) [7]. In males, both a head horn and a thoracic horn become slimmer during pupal remodeling, whereas in females, substantial portion of the pupal head horn disappears to form small three protrusions (Fig. S1). Thus, we also investigated whether such sexually dimorphic remodeling during pupal periods within horn primordia were also regulated by *Tdic-dsx*. We injected dsRNA targeting *Tdic-dsx* into male larvae at several developmental stages (Table S3). We found that adult males treated with *Tdic-dsx* dsRNA at later stages (after 13 h before pupal-chamber formation (BPF)) formed thickened horns similar to a pupal thoracic horn before remodeling (Fig. 5E), whereas the head horn would be properly formed (Fig. S2). These data indicate that DsxM is required for horn remodeling only in the thorax, and is dispensable in the head horn remodeling. On the other hand, when we injected *Tdic-dsx* dsRNA at earlier stages (before 85 h BPF), males did not form adult thoracic horns although small and short thoracic horns were formed during pupal stage (Fig. 5F). These data indicate that DsxM is also required for prepupal and pupal horn growth both in the head and the thoracic horns.

Finally, we also tested whether *Tdic-dsx* is upregulated region-specifically in a sexually dimorphic head horn primordum as reported in the sexually dimorphic structure formation in other insects [17-21]. We performed qRT-PCR at 36 h APF, and found that *Tdic-dsxM* actually showed significantly higher expression in head primordium epidermis than in the surrounding head epidermis (Fig. 5G, H). On the other hand, *Tdic-dsxF* did not show significantly higher expression in the horn primordium at this stage.

## Discussion

Here, we have described development of sexually dimorphic horn in a Japanese rhinoceros beetle, *T. dichotomus*, from both morphological and genetic points of view. We focused on *Tdic-dsx* as the major regulatory gene for sexually dimorphic horn formation and investigated its time of action by knocking down sex determination genes in multiple developmental timepoints during horn morphogenesis period through the use of RNAi. To manipulate sex-specific alternative splicing of *Tdic-dsx*, we identified *Tdic-tra* as a key regulatory factor for the sex-specific splicing of *Tdic-dsx*. We discuss below the genetic regulatory framework of sex determination in *T. dichotomus*, the detailed actions of *Tdic-dsx* in males and females to form sexually dimorphic horns, and evolutionary background realizing the exaggerated horn formation in horned beetles.

### Genetic regulatory mechanisms of sex determination in *T. dichotomus*

Previous studies focusing on horned beetle revealed that *dsxM* promotes male-specific traits including exaggerated horn formation whereas *dsxF* promotes female-specific traits including suppression of horn formation (*Trypoxylus*; [8], *Onthophagus*; [14]). Our RNAi experiments targeting the orthologs of *D. melanogaster* sex determination genes revealed that the sex-specific splicing generating *dsxM* and *dsxF* is regulated by *Tdic-tra* and *Tdic-tra2*, but not by *Tdic-Sxl* during larval-pupal development (Fig. 3C).

Concerning Tdic-Tra and Tdic-Tra2, their function as splicing factors targeting *Tdic-dsx* pre-mRNA seems to be conserved as in other holometabolous insects (e.g. *D. melanogaster,* the housefly *M. domestica, C. capitata, L. cuprina*, *N. vitripennis, B. dorsalis* and *T. castaneum*) [37-44]. The regulatory mechanisms of the sex-specific splicing of *dsx* may be similar to those in *D. melanogaster*. The RNA binding motif (RRM) is present only in Tra2 (Fig. 2A), and the conserved domain structures necessary for target mRNA recognition also support this hypothesis [22,23]. In addition to the molecular functions, biological functions of these genes are at least similar to those in other beetles. The RNAi phenotypes of *tra* and *tra2* orthologs (i.e. the viable masculinized phenotype in *Tdic-tra* RNAi females, and the lethal phenotype in *Tdic-tra2* RNAi females) were comparable with those reported in other beetles (*T. castaneum*; [44], the stag beetle *Cyclommatus metallifer*; [45]). Therefore, the function of Tra and Tra2 as sex determination factors targeting *dsx,* and the function of Tra2 as an essential factor progressing normal development during larval-pupal development may be conserved among Polyphaga, the coleopteran suborder including the above three beetles.

In *D. melanogaster,* Ix directly binds to DsxF but not to DsxM, and function as a co-activator to facilitate transcription activity of target genes of DsxF [32]. The intersexual phenotype observed only in *Tdic-ix* RNAi-treated females (Fig. 3A) implies that Tdic-Ix might also interact with Tdic-DsxF to regulate female-specific sex differentiation as in *D. melanogaster*. Although we did not investigate the detailed molecular interaction between Ix and DsxF in *T. dichotomus*, a female-specific intersexual transformation phenotype reported in the stag beetle *C. metallifer* [45] suggests that the female-specific function of *ix* may be conserved among Polyphaga as well.

Contrasting with the above functionally conserved genes, our RNAi experiments implied that Tdic-Sxl did not regulate the sex-specific alternative splicing of *Tdic-tra*, which is a direct regulatory target of Sxl in *D. melanogaster*. In fact, splicing of *tra* orthologs are not regulated by *Sxl* orthologs even in other Dipteran species [46,47]. In holometabolous insects so far except for *Drosophila* species, the sex-specific alternative splicing of *tra* pre-mRNA is performed through the contribution of the TRA own protein [48,49]. Thus, the sex-specific alternative splicing of *Tdic-tra* would also be regulated by Tdic-TRA own protein or unknown species-specific genes other than *Tdic-Sxl*.

To summarize the regulatory mechanisms of *T. dichotomus* sex determination discussed above, functional Tdic-Tra is first expressed female-specifically through species-specific unknown mechanisms. Then, a Tdic-Tra/Tdic-Tra2 heterodimer would be formed only in females and produce *Tdic-ds*x*F* mRNA and Tdic-DsxF to promote female differentiation by interacting with Tdic-Ix (Fig. 6). On the other hand, in males, functional Tdic-Tra is not expressed, and default splicing of *Tdic-dsx* would result in *Tdic-dsxM* mRNA and Tdic-DsxM production to promote male differentiation. This genetic regulation, in which Tra/Tra2 controls the sex-specific alternative splicing of *dsx*, is shared with other holometabolous insects investigated so far, except for some holometabolous insects in which *tra* ortholog is lost [48]. Therefore, *T. dichotomus* seems to also utilize conserved core regulatory mechanisms shared among holometabolous insects to regulate sex-specific splicing of *dsx* and expression of *dsx*-mediated sexually dimorphic traits.

**Fig. 6.**
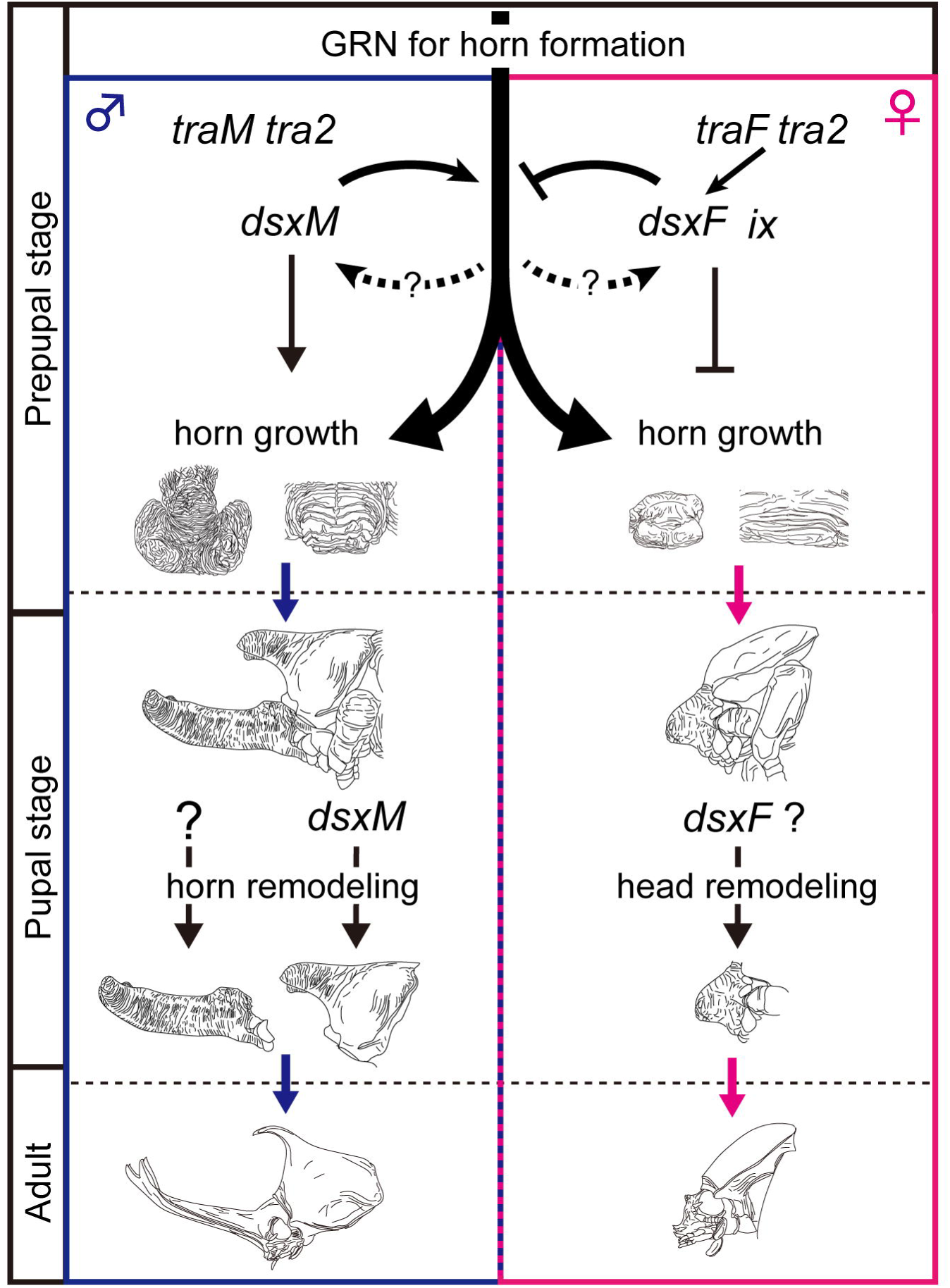
A regulatory model for the formation of sexually dimorphic horns in *T. dichotomus*. In males, default splicing generates *Tdic-dsxM*, whereas in females, *Tdic-tra* (a female isoform, *traF*) and *Tdic-tra2*, generates *Tdic-dsxF* by female-specific splicing of *Tdic-dsx* pre-mRNA. Prepupal *Tdic-dsxM* expression in males positively regulates the growth of head and thoracic horn primordia, whereas prepupal *Tdic-dsxF* in females suppresses the horn growth activity. In pupa, *Tdic-dsxM* regulates thoracic horn remodeling, but is dispensable for remodeling of head horn. GRN, gene regulatory network.

### The gene regulatory network driving horn dimorphism formation in *T. dichotomus*

A previous study revealed that *Tdic-dsx* RNAi in both males and females does not result in a total loss of horns, but results in intermediate-lengthened head horn formation and loss of a thoracic horn (Fig. 3) [8]. The head horn phenotype suggests that there exists a GRN for head horn formation that is operated independently of *Tdic-dsx*. *Tdic-dsxM* seems to enhance this GRN whereas *Tdic-dsxF* seems to suppress it. On the other hand, the thoracic horn phenotypes suggest that thoracic horn formation seems to be totally dependent on *Tdic-dsxM* function, and that action of *Tdic-dsxM* on horn formation might be different between head and thoracic horns during prepupa and/or pupa. However, in the course of our *Tdic-dsx* RNAi experiments, we noticed that thoraces of severely intersexually transformed individuals are always slightly bulged at the thoracic horn formation region (Fig. 3A). This implies that *Tdic-dsxF* expression in the female thorax still has suppressive activity against the GRN for thoracic horn formation. If this is the case, the regulatory relation between *Tdic-dsx* and GRN for horn formation has partial similarity between the head and the thorax. The phenotypic difference previously described in head and thoracic horns would be due to difference in length of horn formed in each body region. In the sections below, we mainly discuss the GRN for horn formation based on our head horn data in which interpretation of phenotypes is less ambiguous.

Our knockdown experiment of *Tdic-tra* revealed that the onset of *Tdic-dsx* modulating GRN for head horn formation is as early as 29 h APF. This is approximately 7 hour before the appearance of sexual dimorphism of head horn primordium. Therefore, *Tdic-dsx* seems to modulate head horn primodium formation just before the onset of tissue growth, and following tissue morphogenesis during prepupal stages.

Several genes involved in the GRN for horn formation have been identified in horned beetle species other than *T. dichotomus*, primarily by focusing on *Drosophila* appendage patterning genes [50,51]. These studies propose an attractive evolutionary model in which large portions of the GRN for proximodistal appendage patterning were recruited to acquire beetle horns in the head and thoracic regions. Still, the overall framework of GRN for horn formation including the key regulatory gene sets involved, spatiotemporal expression dynamics, and regulatory relation among those genes, has not been unveiled so far. Therefore, investigating regulatory relations between appendage patterning genes and other head/thoracic patterning genes expressed in horn primordia focusing on this developmental timepoint will lead to verification of this model in the future studies. In addition, because candidates of the *dsx* target genes involved in GRN for horn formation were identified recently in a horned beetle, *O. taurus* [52], functional analyses focusing on orthologs of those genes expressed in *Trypoxylus* horn primordia at this timepoint will also lead to understanding of conserved and divergent aspects of sexually dimorphic horn formation in horned beetles.

Region-specific upregulation of *dsx* is another essential feature to drive GRN for sexual dimorphism formation in insects [17-20]. Intense studies focusing on regulatory mechanisms of sex comb formation in *D. melanogaster* revealed that such region-specific upregulation of *dsx* is mediated by a positive feedback loop between *dsx* and a Hox gene, *Sex combs reduced* [18]. Therefore, region-specific upregulation of *Tdic-dsx* observed in a head horn primordium might reflect an analogous positive feedback loop mechanism. Future research focusing on the patterning mechanisms of beetle horns and its interaction with *dsx* will be another important issue to discuss conserved genetic framework for sexual dimorphism formation.

*Tdic-dsx-*mediated sexually dimorphic morphogenesis in head and thoracic horn formation during larval-pupal development

As previously reported, *Tdic-dsx* RNAi from early stages (Fig. 3A) resulted in short horn formation in males and females. These data indicate that during the prepupal stage *Tdic-dsxM* promotes the growth of head and thoracic horn primordium, whereas *Tdic-dsxF* suppresses growth of head and thoracic horn primordium (Fig. 6, prepupal stage) [8]. On the other hand, RNAi treatments in later stages revealed distinct functions of *Tdic-dsx* in sexually dimorphic horn formation. During pupal-adult development, *Tdic-dsxM* regulates remodeling of a thoracic horn primordium from a rounded shape to a slender hooked shape (Fig. 5E, F, Fig. 6, pupal stage), but is dispensable for head horn remodeling (Fig. S2). These results indicate that *Tdic-dsxM* and *Tdic-dsxF* regulate different aspects of morphogenesis at both prepupal and pupal stages. Furthermore, to what extent *Tdic-dsx* is required during larval-pupal development is also finely regulated during male head formation, male thoracic horn formation, female head protrusion formation, and female flat prothorax formation. Importantly, in contrast to the previous study that suggested that functions of *Tdic-dsxM* and *Tdic-dsxF* in *T. dichotomus* head and thoracic horn formation are to some extent analogous as described in the previous section (that is, in both of the head and the thoracic horn formation, *Tdic-dsxM* functions as a positive regulator whereas *Tdic-dsxF* functions as a negative regulator) [8], our experiments clarified that both spatial cues (i.e. different developmental contexts in head and prothorax) and temporal cues (i.e. different developmental contexts in prepupa and pupa) modulate the actions of *Tdic-dsxM* and *Tdic-dsxF* to drive appropriate morphogenetic activity in each horn primordium at each developmental stage.

A notable feature of *Tdic-dsx* during sexually dimorphic development is that the onset of action during horn development seems to be categorized earlier (i.e. during the prepupal period) than that of many other sexually dimorphic adult traits in holometabolous insects reported so far (i.e. the pupal period). In *D. melanogaster* sex comb formation, *Papilio* mimetic wing morph formation, and *Bicyclus* wing pheromone gland formation, tissue-specific high expression of *dsx* orthologs are detected after pupation [18-21]. Such a difference in *dsx*’s onset of action seems to be due to the requirement of drastic growth during sexually dimorphic structure formation. In either case mentioned above, the finally formed structure accompanies little to no tissue-level drastic growth during development. On the other hand, as in *T. dichotomus* horn formation, *D. melanogaster* genital organ formation, and *C. metallifer* stag beetle mandible formation, whose sexually dimorphic morphogenesis is regulated by *dsx*, accompanies drastic sexually dimorphic growth during larval-pupal development [17, 53-55], and its onset of action is as early as the prepupal stage. Association between *dsx*’s earlier onset of action and requirement of drastic growth during sexual dimorphism formation suggest that recruitment of the *dsx* function in the earlier stage may be one of the prerequisites to form structurally drastically different sexual dimorphism in insects.

The intermediate head horn formation via *Tdic-tra* RNAi in females (Fig. 5A-C) revealed that the anterior region of male head horn and female small protrusions are not homologous and are instead formed from different regions in the clypeolabrum. Such a distinct developmental origin of head horn within the clypeolabrum is also clearly demonstrated in *Onthophagus* species of dung beetles [3], in which the *dsx* orthologs regulate the formation of distinct sexually dimorphic horns in either the anterior or the posterior region of the head. Analogy of multiple horn formation regions within the clypeolabrum in distinct horned beetle species implies that clypeolabrum is a hotspot of morphological innovations in horned beetles. A comprehensive understanding of the GRN for horn formation in both *Trypoxylus* and *Onthophagus* and comparative developmental studies in the future will lead to understanding of molecular mechanisms underlying the evolutionary origin and evolvability of exaggerated horns in beetles.

Here we described the accurate developmental time course of horn primordial morphogenesis during the prepupal stage in the horned beetle *T. dichotomus* using a time-lapse photography system. In addition, we functionally characterized both *Tdic-tra* and *Tdic-tra2,* genes that regulate the sex-specific splicing of *Tdic-dsx*. By manipulating expression levels of *Tdic-tra* and *Tdic-dsx* during different developmental time points, and by quantifying the extent of sex transformation, we revealed the following three crucial features of *Tdic-dsx* function during the development of sexually dimorphic horn formation: (1) *Tdic-dsx* modulates the GRN for horn formation as early as 29 h APF, a timepoint which corresponds to 7 h before sexual dimorphisms of horn primordia first appears; (2) *Tdic-dsx* regulates different aspects of the tissue growth, tissue death and tissue movement of horn primordia depending on both spatial (head/prothorax) and temporal (prepupa/pupa) contexts; (3) *Tdic-dsxM* and *Tdic-dsxF* promotes the formation of outgrowth structure in distinct regions within the clypeolabrum. These findings inform our understanding of the patterning mechanisms at play during *T. dichotomus* horn formation, as well as provide information regarding regulatory shifts in these mechanisms during the evolution of sexually dimorphic traits in horned beetles. The present study provides a good starting point to elucidate such issues.

## Materials and Methods

### Insects

We purchased *T. dichotomus* larvae from Loiinne (Japan), and Urakiso Tennen Kabuto no Sato (Japan). The last instar larvae were sexed as described previously [8], individually fed on humus in plastic containers, and kept at 10 °C until use. Larvae were moved to room temperature at least 10 days, and reared at 28 °C.

### Micro-CT analysis

A male head tissue at 24 hours after pupal-chamber formation (24 h APF) was fixed in Carnoy solution at room temperature overnight, washed in 70% ethanol and stored in 70% ethanol. The sample was rehydrated through a graded ethanol series, and stained with 25% Lugol solution [56-58] for 5 days. The stained sample was scanned using an X-ray micro-CT device (ScanXmate-E090S105, Comscantechno Co., Ltd., Japan) at a tube voltage peak of 60 kVp and a tube current of 100 μA. The sample was rotated 360 degrees in steps of 0.24 degrees, generating 1500 projection images of 992 × 992 pixels. The micro-CT data were reconstructed at an isotropic resolution of 13.3 × 13.3 × 13.3 μm, and converted into an 8-bit tiff image dataset using coneCTexpress software (Comscantechno Co., Ltd., Japan). Three-dimensional tomographic images were obtained using the OsiriX MD software (version 9.0, Pixmeo, SARL, Switzerland) and Imaris software (version 9.1, Carl Zeiss Microscopy Co., Ltd., Japan). Supplemental videos were edited using Adobe Premiere Pro CC (Adobe Systems Co., Ltd., Japan).

### Time-lapse photography

To monitor the precise time course of the morphogenetic changes of male and 16 female horn primordia during larval-pupal development, time-lapse photography was performed at 28 °C every 30 minutes until they had developed into adults using a CMOS camera (VCC-HD3300, SANYO, Co., Japan).

### Observation of horn primordia development based on stable prepupal staging

We found that the head-rocking behavior observed at the end of pupal-chamber formation can be an unambiguous developmental marker for initiation of the prepupal stage (0 h APF). Using this marker, sampling of horn primordia for staging was performed every 12 hours until they had developed into pupae (120 h APF) using time-lapse photography. Head and thoracic horn primordia were dissected out from larvae in 0.75% sodium chloride, and fixed for 90 minutes at room temperature with 4% paraformaldehyde in phosphate buffered saline (PBS). After being washed twice in PBS, photographic images were obtained with a digital microscope (VHX-900, KEYENCE, Co., Japan).

### BLAST search for sex determination genes

We searched for orthologs of the sex determination genes in the *T. dichotomus* RNA-seq database (Accession number) using full-length complementary DNA (cDNA) sequences of *D. melanogaster* genes (*Sxl*, *tra*, *tra2*, *ix*) as query sequences using the tblastn program (https://blast.ncbi.nlm.nih.gov/Blast.cgi). We evaluated orthology of those genes by performing reciprocal tblastx searches against the *D. melanogaster* cDNA database (r6.06) using the identified *T. dichotomos* genes as queries. We deposited cloned partial cDNA sequences of *T. dichotomus* genes at the DNA Data Bank of Japan (DDBJ)/European Molecular Biology Laboratory (EMBL)/GenBank Accession number for *Tdic-Sxl* is LC385009, for *Tdic-tra* are LC385010 - LC385012, for *Tdic-tra2* are LC385013 - LC385018, and for *Tdic-ix* is LC385019.

### cDNA library construction, reverse transcription-polymerase chain reaction (RT-PCR), and sequencing

Total RNA was extracted from each of head or thoracic horn primordia in wild type and RNAi-treated individuals (*EGFP*, *Sxl*, *tra*, *tra2*, *ix, dsx* and *dsxF*) using TRI Reagent (Molecular Research Center, Inc., USA) according to the manufacturer’s instructions. First-stranded cDNA was synthesized with the SuperScript III Reverse Transcriptase (Life Technologies Japan Ltd., Japan) using 1 μg total RNA as a template. Primer sets for cloning and double stranded RNA (dsRNA) synthesis were designed based on the cDNA sequences identified above (Table S1). PCR was performed using Ex Taq DNA polymerase (Takara Bio Inc., Japan) according to the manufacturer’s protocol. Amplified PCR products were purified using MagExtractor (TOYOBO, Co., Ltd., Japan), and subcloned into the pCR4-TOPO vector using the TOPO TA cloning Kit (Life Technologies Japan Ltd., Japan). Sequences of the inserts were determined by a DNA sequencing service at FASMAC Co. Ltd., Japan.

### Gene expression analysis by quantitative RT-PCR (qRT-PCR)

Primer sets for qRT-PCR were designed using Primer3Plus program (http://primer3plus.com/cgi-bin/dev/primer3plus.cgi) (Table S1). qRT-PCR was performed using the cDNA libraries synthesized above and THUNDERBIRD SYBR qPCR Mix (TOYOBO, Co., Ltd., Japan) according to the manufacturer’s instructions. The relative quantification in gene expression was determined using the 2^-ΔΔCt^ method [59].

### Larval RNAi

Primer pairs for dsRNA synthesis were designed within the common regions shared by male and female isoforms (Fig. 2A, Table S1). Partial sequences of the target sequences were amplified by PCR using primers flanked with the T7 promoter sequence in the 5’-ends. DsRNAs were synthesized using AmpliScribe T7-Flash Transcription Kit (Epicentre Technologies, Corp., USA). The purified PCR products were used as templates. Injections of dsRNA were performed as described previously [8]. DsRNA was injected into each late last instar larva or prepupa under the conditions described in Table S2 and Table S3. *Enhanced green fluorescent protein* (*EGFP*) dsRNA was injected as a negative control.

### Horn and body length measurement

Horn length was measured by extracting the contour of a horn in the lateral view using the SegmentMeasure plugin for ImageJ 64 developed by Hosei Wada. We defined a body length as the length between the anterior tip of the clypeus in the head to the posterior most region of the abdomen, and was directly measured using a digital caliper (DN-100, Niigata seiki, Co., Ltd., Japan)

## Acknowledgments

We thank Hosei Wada for developing the “SegmentMeasure” plug-in of ImageJ, Takahiro Ohde, Hiroki Gotoh, and Yasuhiko Chikami for helpful discussion, Robert Zinna for critical reading of the manuscript, the Model Plant Research Facility/NIBB BioResource Center and the Functional Genomics Facility/NIBB Core Research Facilities for technical assistance. TN was supported by MEXT KAKENHI (23128505, 25128706, 16H01452 and 18H04766) and NIBB Cooperative Research Programs (18-433).

## Movie 1 Micro-CT analysis of a male prepupal head at 24 h APF.

(a) Head horn primordium formation regions within clypeolabrum. A small epidermal protrusion in the clypeolabral, which seems to be a head horn primordium, was observed above a clypeal primordium (Fig. 1A). The head capsule is indicated in blue [60]. (b) The morphological character of the ocular and the mandibular apodeme at 24 h APF. Apolysis was incomplete in the ocular and the mandibular apodeme at 24 h APF [35,36]. After this stage, prepupal horn primordia can be readily dissected out due to apolysis at the ocular and the mandibular apodeme (Fig. 1B). This morphological character is unambiguous developmental marker to know the onset of sexual dimorphism formation (36 h APF).

## Movie 2 Time-lapse photography of last instar larvae.

A head-rocking behavior starts to be observed at the end of pupal chamber formation.

---

**Fig. S1 Morphological comparison of pupal horn primordia and adult horns in *T. dichotomus.***

Pupal horn primordia are rounded, and slightly larger than adult horns. Light blue in pupal horn primordia shows the morphologies of adult male horn or female three small protrusions.

**Fig. S2 Head horn formation phenotypes induced by *Tdic-dsx* RNAi at late prepupal stages.**

Head horn remodeling during pupal-adult development in males. (A, A’) A wild type male head horn. (B, B’) Head horn formed by a late *Tdic-dsx* RNAi treatment (13 BPF). (C, C’) Head horn formed by an early *Tdic-dsx* RNAi treatment (85 BPF). *Tdic-dsxM* is dispensable for head horn remodeling. (A) – (C) the lateral views of a head horn, (A’) – (C’) the dorsal views of a head horn tips. Scale bars are 5 mm.

**Table S1 Primers used in this study.**

**Table S2 RNAi treatment conditions in Fig.3.**

**Table S3 RNAi treatment conditions in Fig.4 and Fig.5.**

## References

1. Eberhard, W.G., 1980. Horned beetles. Sci. Am. 242, 166–183. http://www.jstor.org/stable/24966285

2. Emlen, D.J., Marangelo, J., Ball, B., Cunningham, C.W., 2007. Diversity in the weapons of sexual selection: horn evolution in the beetle genus *Onthophagus* (coleoptera: scarabaeidae). Evolution 59, 1060–1084. https://doi.org/10.1111/j.0014-3820.2005.tb01044.x

3. Busey, H.A., Zattara, E.E., Moczek, A.P., 2016. Conservation, innovation, and bias: Embryonic segment boundaries position posterior, but not anterior, head horns in adult beetles. J. Exp. Zool. B. Mol. Dev. Evol. 326, 271–279. https://doi.org/10.1002/jez.b.22682

4. McCullough, E.L., Tobalske, B.W., Emlen, D.J., 2014. Structural adaptations to diverse fighting styles in sexually selected weapons. Proc. Natl. Acad. Sci. U.S.A 111, 14484–14488. https://doi.org/10.1073/pnas.1409585111

5. Siva-Jothy, M.T., 1987. Mate securing tactics and the cost of fighting in the Japanese horned beetle, *Allomyrina dichotoma* L. (Scarabaeidae). J. Ethol. 5, 165– 172. https://doi.org/10.1007/BF02349949

6. Hongo, Y., 2003. Appraising behaviour during male-male interaction in the Japanese horned beetle *Trypoxylus dichotomus septentrionalis* (Kono). Behaviour 140, 501–517. https://doi.org/10.1163/156853903322127959

7. Emlen, D.J., Lavine, L.C., Ewen-Campen, B., 2007a. On the origin and evolutionary diversification of beetle horns. Proc. Natl. Acad. Sci. U.S.A. 104 Suppl 1, 8661–8668. https://doi.org/10.1073/pnas.0701209104

8. Ito, Y., Harigai, A., Nakata, M., Hosoya, T., Araya, K., Oba, Y., Ito, A., Ohde, T., Yaginuma, T., Niimi, T., 2013. The role of *doublesex* in the evolution of exaggerated horns in the Japanese rhinoceros beetle. EMBO Rep. 14, 561–567. https://doi.org/10.1038/embor.2013.50

9. Matsuda, K., Gotoh, H., Tajika, Y., Sushida, T., Aonuma, H., Niimi, T., Akiyama, M., Inoue, Y., Kondo, S., 2017. Complex furrows in a 2D epithelial sheet code the 3D structure of a beetle horn. Sci. Rep. 7, 13939. https://doi.org/10.1038/s41598-017-14170-w

10. Moczek, A.P., Cruickshank, T.E., Shelby, A., 2006. When ontogeny reveals what phylogeny hides: gain and loss of horns during development and evolution of horned beetles. Evolution 60, 2329. https://doi.org/10.1554/06-260.1

11. Moczek, A.P., 2007. Pupal remodeling and the evolution and development of alternative male morphologies in horned beetles. BMC Evol. Biol. 7, 151. https://doi.org/10.1186/1471-2148-7-151

12. Cline, T.W., Meyer, B.J., 1996. Vive la différence: males vs females in flies vs worms. Annu. Rev. Genet. 30, 637–702. https://doi.org/10.1146/annurev.genet.30.1.637

13. Hediger, M., Burghardt, G., Siegenthaler, C., Buser, N., Hilfiker-Kleiner, D., Dübendorfer, A., Bopp, D., 2004. Sex determination in *Drosophila melanogaster* and *Musca domestica* converges at the level of the terminal regulator *doublesex*. Dev. Genes. Evol. 214, 29–42. https://doi.org/10.1007/s00427-003-0372-2

14. Kijimoto, T., Moczek, A.P., Andrews, J., 2012. Diversification of *doublesex* function underlies morph-, sex-, and species-specific development of beetle horns. Proc. Natl. Acad. Sci. U.S.A. 109, 20526–20531. https://doi.org/10.1073/pnas.1118589109

15. Shukla, J.N., Palli, S.R., 2012. Doublesex target genes in the red flour beetle, *Tribolium castaneum*. Sci. Rep. 2, 948. https://doi.org/10.1038/srep00948

16. Suzuki, M.G., Funaguma S., Kanda T., Tamura T., Shimada T., 2005. Role of the male BmDsx protein in the sexual differentiation of *Bombyx mori*. Evol. Dev. 7, 58–68. https://doi.org/10.1111/j.1525-142X.2005.05007.x

17. Gotoh, H., Miyakawa, H., Ishikawa, A., Ishikawa, Y., Sugime, Y., Emlen, D.J., Lavine, L.C., Miura, T., 2014. Developmental link between sex and nutrition; *doublesex* regulates sex-specific mandible growth via juvenile hormone signaling in stag beetles. PLOS Genet. 10, e1004098. https://doi.org/10.1371/journal.pgen.1004098

18. Tanaka, K., Barmina, O., Sanders, L.E., Arbeitman, M.N., Kopp, A., 2011. Evolution of sex-specific traits through changes in HOX-dependent *doublesex* expression. PLoS Biol. 9, e1001131. https://doi.org/10.1371/journal.pbio.1001131

19. Rice, G., Barmina, O., Hu, K., Kopp, A., 2018. Evolving *doublesex* expression correlates with the origin and diversification of male sexual ornaments in the *Drosophila* immigrans species group. Evol. Dev. 20, 78–88. https://doi.org/10.1111/ede.12249

20. Kunte, K., Zhang, W., Tenger-Trolander, A., Palmer, D.H., Martin, A., Reed, R.D., Mullen, S.P., Kronforst, M.R., 2014. *doublesex* is a mimicry supergene. Nature 507, 229–232. https://doi.org/10.1038/nature13112

21. Bhardwaj, S., Prudic, K.L., Bear, A., Dasgupta, M., Wasik, B.R., Tong, X., Cheong, W.F., Wenk, M.R., Monteiro, A., 2018. Sex differences in 20-hydroxyecdysone hormone levels control sexual dimorphism in *Bicyclus anynana* wing patterns. Mol. Biol. Evol. 35, 465–472. https://doi.org/10.1093/molbev/msx301

22. Schütt, C., Nöthiger, R., 2000. Structure, function and evolution of sex-determining systems in Dipteran insects. Development 127, 667–677. http://dev.biologists.org/content/127/4/667.long

23. Bell, L.R., Maine, E.M., Schedl, P., Cline, T.W., 1988. *Sex-lethal*, a Drosophila sex determination switch gene, exhibits sex-specific RNA splicing and sequence similarity to RNA binding proteins. Cell 55, 1037–1046. https://doi.org/10.1016/0092-8674(88)90248-6

24. Samuels, M.E., Bopp, D., Colvin, R.A., Roscigno, R.F., Garcia-Blanco, M.A., Schedl, P., 1994. RNA binding by Sxl proteins in vitro and in vivo. Mol. Cell. Biol. 14, 4975–4990. https://doi.org/10.1128/MCB.14.7.4975

25. Samuels, M., Deshpande, G., Schedl, P., 1998. Activities of the Sex-lethal protein in RNA binding and protein:protein interactions. Nucl. Acids Res. 26, 2625–2637. https://doi.org/10.1093/nar/26.11.2625

26. Inoue, K., Hoshijima, K., Sakamoto, H., Shimura, Y., 1990. Binding of the *Drosophila Sex-lethal* gene product to the alternative splice site of *transformer* primary transcript. Nature 344, 461–463. https://doi.org/10.1038/344461a0

27. Inoue, K., Hoshijima, K., Higuchi, I., Sakamoto, H., Shimura, Y., 1992. Binding of the *Drosophila transformer* and *transformer-2* proteins to the regulatory elements of *doublesex* primary transcript for sex-specific RNA processing. Proc. Natl. Acad. Sci. U.S.A. 89, 8092–8096. https://doi.org/10.1073/pnas.89.17.8092

28. Amrein, H., Hedley, M.L., Maniatis, T., 1994. The role of specific protein-RNA and protein-protein interactions in positive and negative control of pre-mRNA splicing by *Transformer 2*. Cell 76, 735–746. https://doi.org/10.1016/0092-8674(94)90512-6

29. Erdman, S.E., Chen, H.J., Burtis, K.C., 1996. Functional and genetic characterization of the oligomerization and DNA binding properties of the *Drosophila doublesex* proteins. Genetics 144, 1639–1652. http://www.genetics.org/content/144/4/1639.long

30. Luo, S.D., Shi, G.W., Baker, B.S., 2011. Direct targets of the *D. melanogaster* DsxF protein and the evolution of sexual development. Development 138, 2761–2771. https://doi.org/10.1242/dev.065227

31. Verhulst, E.C., van de Zande, L., 2015. Double nexus—Doublesex is the connecting element in sex determination. Brief. Funct. Genomics 14, 396–406. https://doi.org/10.1093/bfgp/elv005

32. Garrett-Engele, C.M., Siegal, M.L., Manoli, D.S., Williams, B.C., Li, H., Baker, B.S., 2002. *intersex*, a gene required for female sexual development in *Drosophila*, is expressed in both sexes and functions together with *doublesex* to regulate terminal differentiation. Development 129, 4661–4675. http://dev.biologists.org/content/129/20/4661.long

33. Posnien, N., Schinko, J.B., Kittelmann, S., Bucher, G., 2010. Genetics, development and composition of the insect head‐‐a beetle’s view. Arthropod Struct. Dev. 39, 399–410. https://doi.org/10.1016/j.asd.2010.08.002

34. Posnien, N., Koniszewski, N.D.B., Hein, H.J., Bucher, G., 2011. Candidate gene screen in the red flour beetle *Tribolium* reveals *six3* as ancient regulator of anterior median head and central complex development. PLoS Genet. 7. https://doi.org/10.1371/journal.pgen.1002416

35. Manton, S.M., Harding, J.P., 1964. Mandibular mechanisms and evolution of arthropods. Phil. Trans. R. Soc. Lond. B 247, 1–183. https://doi.org/10.1098/rstb.1964.0001

36. Soler, C., Laddada, L., Jagla, K., 2016. Coordinated development of muscles and tendon-like structures: early interactions in the *Drosophila* leg. Front. Physiol. 7, 22. https://doi.org/10.3389/fphys.2016.00022

37. Hoshijima, K., Inoue, K., Higuchi, I., Sakamoto, H., Shimura, Y., 1991. Control of *doublesex* alternative splicing by *transformer* and *transformer-2* in *Drosophila*. Science 252, 833–836. https://doi.org/10.1126/science.1902987

38. Hediger, M., Henggeler, C., Meier, N., Perez, R., Saccone, G., Bopp, D., 2010. Molecular characterization of the key switch *F* provides a basis for understanding the rapid divergence of the sex-determining pathway in the housefly. Genetics 184, 155–170. https://doi.org/10.1534/genetics.109.109249

39. Pane, A., De Simone, A., Saccone, G., Polito, C., 2005. Evolutionary conservation of *Ceratitis capitata transformer* gene function. Genetics 171, 615–624. https://doi.org/10.1534/genetics.105.041004

40. Concha, C., Scott, M.J., 2009. Sexual development in *Lucilia cuprina* (Diptera, Calliphoridae) is controlled by the transformer gene. Genetics 182, 785–798. https://doi.org/10.1534/genetics.109.100982

41. Verhulst, E.C., Beukeboom, L.W., van de Zande, L., 2010. Maternal control of haplodiploid sex determination in the wasp *Nasonia*. Science 328, 620–623. https://doi.org/10.1126/science.1185805

42. Liu, G., Wu, Q., Li, J., Zhang, G., Wan, F., 2015. RNAi-mediated knock-down of *transformer* and t*ransformer 2* to generate male-only progeny in the oriental fruit fly, *Bactrocera dorsalis* (Hendel). PLOS ONE 10, e0128892. https://doi.org/10.1371/journal.pone.0128892

43. Shukla, J.N., Palli, S.R., 2012. Sex determination in beetles: production of all male progeny by parental RNAi knockdown of *transformer*. Sci. Rep. 2, 602. https://doi.org/10.1038/srep00602

44. Shukla, J.N., Palli, S.R., 2013. *Tribolium castaneum* Transformer-2 regulates sex determination and development in both males and females. Insect Biochem. Mol. Biol. 43, 1125–1132. https://doi.org/10.1016/j.ibmb.2013.08.010

45. Gotoh, H., Zinna, R.A., Warren, I., DeNieu, M., Niimi, T., Dworkin, I., Emlen, D.J., Miura, T., Lavine, L.C., 2016. Identification and functional analyses of sex determination genes in the sexually dimorphic stag beetle *Cyclommatus metallifer*. BMC Genomics 17, 250. https://doi.org/10.1186/s12864-016-2522-8

46. Meise, M., Hilfiker-Kleiner, D., Dübendorfer, A., Brunner, C., Nöthiger, R., Bopp, D., 1998. *Sex-lethal*, the master sex-determining gene in *Drosophila*, is not sex-specifically regulated in *Musca domestica*. Development 125, 1487–1494. http://dev.biologists.org/content/125/8/1487.long

47. Saccone, G., Peluso, I., Artiaco, D., Giordano, E., Bopp, D., Polito, L.C., 1998. The *Ceratitis capitata* homologue of the *Drosophila* sex-determining gene *Sex-lethal* is structurally conserved, but not sex-specifically regulated. Development 125, 1495– 1500. http://dev.biologists.org/content/125/8/1495.long

48. Geuverink, E., Beukeboom, L.W., 2014. Phylogenetic distribution and evolutionary dynamics of the sex determination genes *doublesex* and *transformer* in insects. Sex.Dev. 8, 38–49. https://doi.org/10.1159/000357056

49. Tanaka, A., Aoki, F., Suzuki, M.G., 2018. Conserved domains in the transformer protein act complementary to regulate sex-specific splicing of its own pre-mRNA. Sex. Dev. https://doi.org/10.1159/000489444

50. Moczek, A.P., Nagy, L.M., 2005. Diverse developmental mechanisms contribute to different levels of diversity in horned beetles. Evol. Dev. 7, 175–185. https://doi.org/10.1111/j.1525-142X.2005.05020.x

51. Moczek, A.P., Rose, D.J., 2009. Differential recruitment of limb patterning genes during development and diversification of beetle horns. Proc. Natl. Acad. Sci. U.S.A. 106, 8992–8997. https://doi.org/10.1073/pnas.0809668106

52. Ledón-Rettig, C.C., Zattara, E.E., Moczek, A.P., 2017. Asymmetric interactions between *doublesex* and tissue‐ and sex-specific target genes mediate sexual dimorphism in beetles. Nat. Commun. 8, 14593. https://doi.org/10.1038/ncomms14593

53. Sanchez, L., Gorfinkiel, N., Guerrero, I., 2001. Sex determination genes control the development of the *Drosophila* genital disc, modulating the response to Hedgehog, Wingless and Decapentaplegic signals. Development 128, 1033–1043. http://dev.biologists.org/content/128/7/1033.long

54. Keisman, E.L., Christiansen, A.E., Baker, B.S., 2001. The sex determination gene *doublesex* regulates the A/P organizer to direct sex-specific patterns of growth in the *Drosophila* genital imaginal disc. Dev. Cell 1, 215–225. https://doi.org/10.1016/S1534-5807(01)00027-2

55. Ahmad, S.M., Baker, B.S., 2002. Sex-specific deployment of FGF signaling in *Drosophila* recruits mesodermal cells into the male genital imaginal disc. Cell 109, 651–661. https://doi.org/10.1016/S0092-8674(02)00744-4

56. Degenhardt, K., Wright, A.C., Horng, D., Padmanabhan, A., Epstein, J.A., 2010. Rapid 3D phenotyping of cardiovascular development in mouse embryos by micro-CT with iodine staining. Circ. Cardiovasc. Imaging 3, 314–322. https://doi.org/10.1161/CIRCIMAGING.109.918482

57. Metscher, B.D., 2009. MicroCT for developmental biology: a versatile tool for high-contrast 3D imaging at histological resolutions. Dev. Dyn. 238, 632–640. https://doi.org/10.1002/dvdy.21857

58. Metscher, B.D., 2009. MicroCT for comparative morphology: simple staining methods allow high-contrast 3D imaging of diverse non-mineralized animal tissues. BMC Physiol. 9, 11. https://doi.org/10.1186/1472-6793-9-11

59. Livak, K.J., Schmittgen, T.D., 2001. Analysis of relative gene expression data using real-time quantitative PCR and the 2(-Delta Delta C(T)) Method. Methods 25, 402– 408. https://doi.org/10.1006/meth.2001.1262

60. Richter, S., Stein, M., Frase, T., Szucsich, N., 2013. The Arthropod Head, in: Arthropod Biology and Evolution: Molecules, Development, Morphology. 223–240. https://doi.org/10.1007/978-3-642-36160-9_10

